# Molecular Mechanisms of Drug Resistance and Compensation in SARS-CoV-2 Main Protease: The Interplay Between E166 and L50

**DOI:** 10.1101/2025.01.24.634813

**Authors:** Sarah N. Zvornicanin, Ala M. Shaqra, Julia Flynn, Heidi Carias Martinez, Weiping Jia, Stephanie Moquin, Dustin Dovala, Daniel N. Bolon, Nese Kurt Yilmaz, Celia A. Schiffer

**Affiliations:** Department of Biochemistry and Molecular Biotechnology, University of Massachusetts Chan Medical School, Worcester, Massachusetts 01605, USA; Biomedical Research, Novartis, Emeryville, CA 94608, USA

## Abstract

The SARS-CoV-2 main protease (M^pro^) is essential for viral replication, and a primary target for COVID-19 antivirals. Direct-acting antivirals such as nirmatrelvir, the active component of Paxlovid, target the M^pro^ active site to block viral polyprotein cleavage and thus replication. However, drug resistance mutations at the active site residue Glu166 (E166) have emerged in *in vitro* selection studies, raising concerns about the durability of current antiviral strategies. Here, we investigate the molecular basis of drug resistance conferred by E166A and E166V mutations against nirmatrelvir and the related PF-00835231, individually and in combination with the distal mutation L50F. We found that E166 mutations reduce nirmatrelvir potency by up to 3000-fold while preserving substrate cleavage, with catalytic efficiency reduced by only up to 2- fold. This loss of catalytic efficiency was compensated for by the addition of L50F in the double-mutant variants. We have determined three cocrystal structures of the E166 variants (E166A, E166V, and E166V/L50F) bound to PF-00835231. Comparison of these structures with wild-type demonstrated that E166 is crucial for dimerization and for shaping the substrate-binding S1 pocket. Our findings highlight the mutability of E166, a prime site for resistance for inhibitors that leverage direct interactions with this position, and the potential emergence of highly resistant and active variants in combination with the compensatory mutation L50F. These insights support the design of inhibitors that target conserved protease features and avoid E166 side-chain interactions to minimize susceptibility to resistance.

## Introduction

The main protease (M^pro^) of SARS-CoV-2 is essential to the viral life cycle and has been a target for small molecule drug development for treatment of COVID-19. Paxlovid, composed of the direct-acting antiviral nirmatrelvir (PF-07321332; a reversible covalent inhibitor of M^pro^) and ritonavir, remains the primary treatment for moderate cases of COVID-19 in the US. Various other M^pro^ inhibitors, including the nirmatrelvir analog PF-00835231^1,2^, have been explored in preclinical or clinical evaluations. Viral protease inhibitors have historically been susceptible to drug resistance mutations. Since the US FDA issued an Emergency Use Authorization for Paxlovid in 2021, drug resistant M^pro^ variants have been observed in *in vitro* resistance selection experiments^3–5^. In addition, the E166V mutation was observed in three Paxlovid-treated patients in the EPIC-HR trial, though live virus was not recovered from these samples, and none of these patients experienced hospitalization or death^5^.

Residue 166 of M^pro^, although highly evolutionarily conserved in coronaviruses, is proposed to be one of the critical sites for drug resistance in SARS-CoV-2 M^pro6,7^, with E166 variants showing the largest decreases in potency against nirmatrelvir^6^. Mutations at this position have been observed in two of the three nirmatrelvir *in vitro* selection studies performed using SARS-CoV-2^3–5^. In agreement with this, we have previously found that evolutionary conservation of a position does not correlate with susceptibility to resistance mutations, and that E166 can tolerate a variety of substitutions^8,9^. E166 variants including E166A/G/H/I/K/L/V/Y have been reported to cause a small to moderate reduction in catalytic efficiency (k_cat_/K_M_) while conferring high levels of nirmatrelvir resistance^6^. The molecular mechanisms underlying drug resistance caused by these mutations remain elusive.

Viral passaging^3,4,10,11^, *in vitro*^6,8,12,13^, and *in silico*^14^ experiments have specifically highlighted nirmatrelvir-resistant M^pro^ variants E166A and E166V. Although these two variant enzymes are highly drug resistant, their activities are reported to be extremely deficient, which may explain their low prevalence in patient populations^4,6^. In SARS-CoV-2 M^pro^, resistance mutations such as E166V have resulted in an adverse effect on the fitness of the virus^3,4^. In other drug resistance systems such as HIV-1 protease, mutations often accumulate in complex combinations to reach significant levels of resistance while retaining fitness^15^. Primary mutations close to the active site typically weaken inhibitor binding, causing drug resistance. Mutations distal to the active site can either contribute to resistance by weakening inhibitor binding or they can facilitate compensation for a variant enzyme’s compromised catalytic function^15^. Many M^pro^ drug resistance mutations are proximal to the inhibitor binding site, including substitutions at E166; in contrast, the non-resistant, compensatory mutation L50F is distal (**Figure 1**). The distal L50F mutation alone results in a hyperactive variant in biochemical assays that does not seem to display resistance against nirmatrelvir^16,17^. L50F has been reported to co-occur with E166 mutations to possibly compensate for the loss in enzymatic activity, and therefore has been suggested as a background mutation that can accelerate resistance selection^3,6,10^. In viral passaging experiments, the particularly compromised E166A and E166V M^pro^ variants evolve the hyperactive and compensatory mutation L50F either shortly before or after the occurrence of the major E166 drug resistance mutation^3,4,10^.

**Figure 1.**
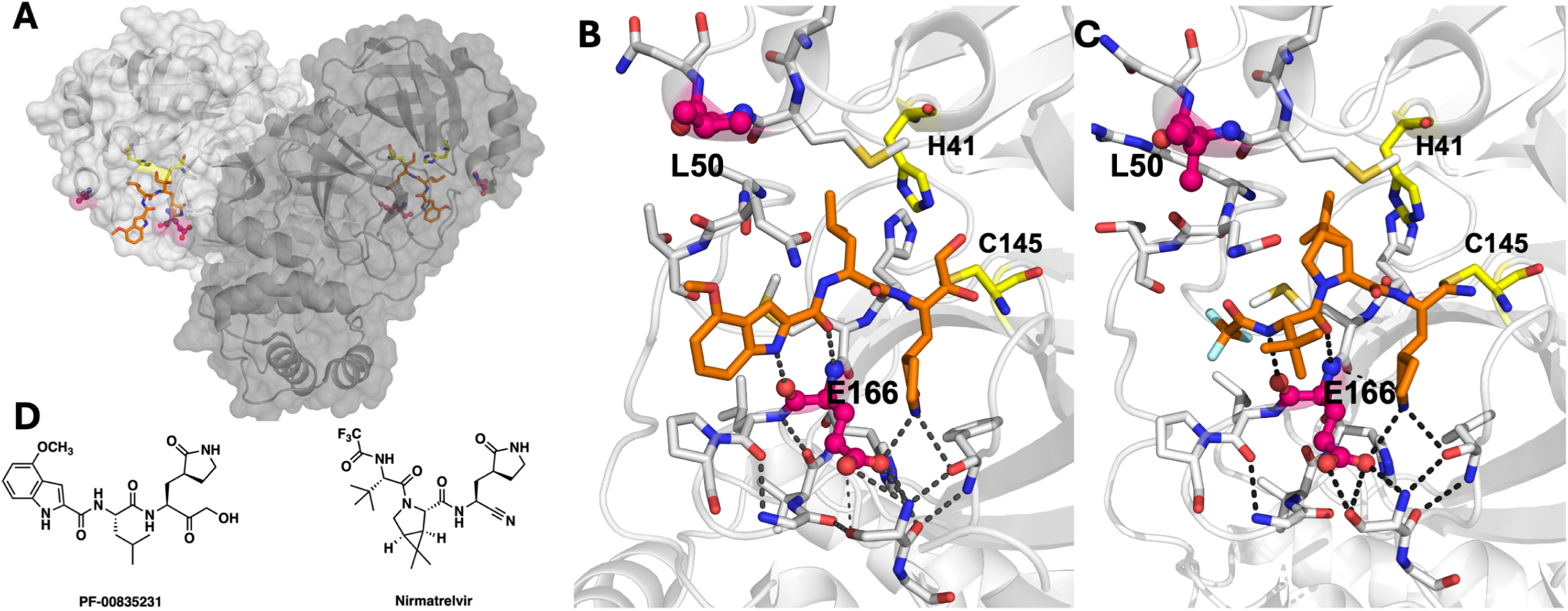
Overview of SARS-CoV-2 M^pro^ structure (PDB ID: 8DSU). **A**) The dimeric M^pro^ crystal structure shown in semi-transparent surface and cartoon representation with each protomer in a shade of gray. **B**) Close-up view of the active site where the dark gray dashed lines represent the hydrogen bonds between PF-00835231, E166, and other residues involved in dimerization. In both panels the catalytic residues (H41 and C145) are in yellow, the two sites of mutation (L50 and E166) are colored magenta, and the inhibitor PF-00835231 covalently bound at the active site is depicted as orange sticks (PDB ID: 8DSU). C) Nirmatrelvir bound to the active site of WT M^pro^ (PDB ID: 7RFS). D) 2D chemical structures of PF- 00835231 and nirmatrelvir.

The combination of the highly drug resistant but catalytically compromised E166A or E166V variants with the compensatory L50F mutation poses a threat against the current COVID-19 direct-acting antivirals, including nirmatrelvir. The mechanism of resistance conferred by E166 mutations, and the role of hyperactivating or compensatory mutations such as L50F in combination with these highly drug-resistant variants need to be explored to counter this threat that can render antivirals ineffective and obsolete. By analyzing the combination of emerging drug-resistant and compensatory variants of SARS-CoV-2 M^pro^, we can better elucidate M^pro^ resistance mechanisms to inform drug design in response to these common mutations. Here, we determined cocrystal structures of E166A and E166V variants of SARS-CoV-2 M^pro^, as well as the double-mutant variant E166V/L50F. To complement these structures, we also assessed single- and double-mutant variant enzyme catalytic efficiencies, protein dimerization, protein thermostability, and inhibition by nirmatrelvir and PF-00835231, all of which we compare to wild-type (WT) M^pro^.

We found that the drug-resistant variants we assessed here are defective in catalytic efficiency and dimerization but are highly resistant to nirmatrelvir. Our structures allowed us to hypothesize that the primary molecular mechanism of resistance at the E166 site is affinity reduction which occurs via the loss of three essential hydrogen bonds made by E166 to shape the essential and conserved P1 glutamine binding pocket at the active site. Our findings also point to L50F as a compensatory mutation, as compounding evidence suggests, even in the background of a highly defective and drug-resistant enzyme variant. Our work thus characterizes the molecular basis for drug resistance against nirmatrelvir and motivates the development of inhibitors that do not rely on interactions with E166 side-chain atoms.

## Results

### Potencies of nirmatrelvir and PF-00835231 are reduced against E166A and E166V M^pro^ variants

To study the impact of mutations at E166, we engineered a series of five M^pro^ variants (L50F, E166A, E166V, E166A/L50F and E166V/L50F). We expressed and purified these enzymes as described in the Methods. To investigate the level of drug resistance they confer, we measured the potencies of both reversibly covalent compounds nirmatrelvir and PF-00835231 (a chemically similar analog from Pfizer that can only be dosed intravenously) against each E166 variant (**Table 1**) using a fluorescence-based enzymatic assay. We quantified drug resistance as the fold-change of inhibitor potency between the WT and variant M^pro^ enzymes.

**Table 1.** Potencies of nirmatrelvir and PF-00835231 against E166 M^pro^ variants. The inhibition constants (*K_i_*) are reported as the mean and SEM from at least three measurements. Fold-change in potency was calculated by dividing the potency of the inhibitor against the variant by the potency against WT M^pro^.

| M<sup>pro</sup> Variant | Nirmatrelvir<br>$K_i$ (μM) ± SEM | Nirmatrelvir<br>Fold-change in<br>Potency | PF-00835231<br>$K_i$ (μM) ± SEM | PF-00835231<br>Fold-change in<br>Potency |
| --- | --- | --- | --- | --- |
| WT | 0.003 ± 0.001 | 1 | 0.0008 ± 0.0003 | 1 |
| L50F | 0.0026 ± 0.0003 | ~1 | ≤ 0.0008 * | ≤ 1 |
| E166A | 0.082 ± 0.006 | 30 | 0.007 ± 0.001 | 10 |
| E166V | 8.2 ± 0.4 | 2700 | $<$ 0.008 * | $<$ 10 |
| E166A/L50F | 0.025 ± 0.003 | 10 | 0.0035 ± 0.0006 | $<$ 10 |
| E166V/L50F | 2.83 ± 0.08 | 950 | 0.010 ± 0.002 | 10 |

\* Inhibition was too potent to measure a reliable $K_i$ and thus reached the assay's limit of detection

In inhibition assays with nirmatrelvir, the E166A variant of SARS-CoV-2 M^pro^ resulted in moderate resistance, with about 30-fold potency loss relative to WT (**Table 1**). The E166V variant resulted in severe nirmatrelvir resistance at a 2700-fold potency decrease relative to WT. E166A/L50F and E166V/L50F variants demonstrated 10- and 950-fold potency loss, respectively. Intriguingly, addition of the L50F mutation to the single-mutant variants resulted in a partial restoration of potency, by about 3-fold for both cases.

In contrast to nirmatrelvir, all variants tested here displayed much lower potency shifts against PF-00835231 (**Table 1**). Each of the variants lost similar, low levels of potency against PF- 00835231, only as much as 10-fold for any variant relative to the WT enzyme.

In summary, we found that the valine variants of M^pro^, E166V and E166V/L50F, showed very large potency losses against nirmatrelvir while potency losses for the analog PF-00835231 were much smaller. Alanine variants had less potency loss for both inhibitors, but still lost between 10- and 30-fold potency relative to WT.

### Enzymatic activity of E166 variants is significantly reduced, and L50F is compensatory

We next tested whether conferring resistance came at the cost of losing enzymatic activity for these variants. The efficiency of cleaving a peptide corresponding to the natural nsp4-nsp5 cleavage site processed by M^pro^ was measured for E166A, E166V, E166A/L50F and E166V/L50F variants. Peptide cleavage was highly compromised across all variants, with less than 40% of the WT catalytic efficiency (k_cat_/K_M_) (**Table 2**, **Figure 2a**). The enzymatic assays revealed that most of the loss in catalytic efficiency resulted from a reduction in k_cat_ rather than a significant change in K_M_, which only varied by at most 2-fold. In contrast, the catalytic efficiency for the E166 variants varied from 6% to 38% compared to WT (**Table 2**). The E166A variant maintained 15% of WT catalytic efficiency, while the E166V variant had only 6%. L50F alone was hyperactive as we observed previously^16^ with 133% of WT catalytic efficiency. Additionally, combination of L50F with the E166 variants partially rescued catalytic activity. When L50F was added to E166V, the double-mutant variant E166V/L50F had increased catalytic activity which was boosted 185% relative to its single-mutant variant, and from just 6% to 11% of WT. The double-mutant variant E166A/L50F also demonstrated an increased catalytic efficiency of 255% relative to its single-mutant variant enzyme, from 15% to 38% of WT. Thus in complement to reports that observed increased viral fitness of E166/L50F mutant viruses compared to E166V mutant viruses^3,4,10^ our biochemical assessment of double-mutant variant E166 enzymes demonstrates that L50F acts as a compensatory mutation that increases enzymatic activity of compromised M^pro^ resistant variants.

**Figure 2:**
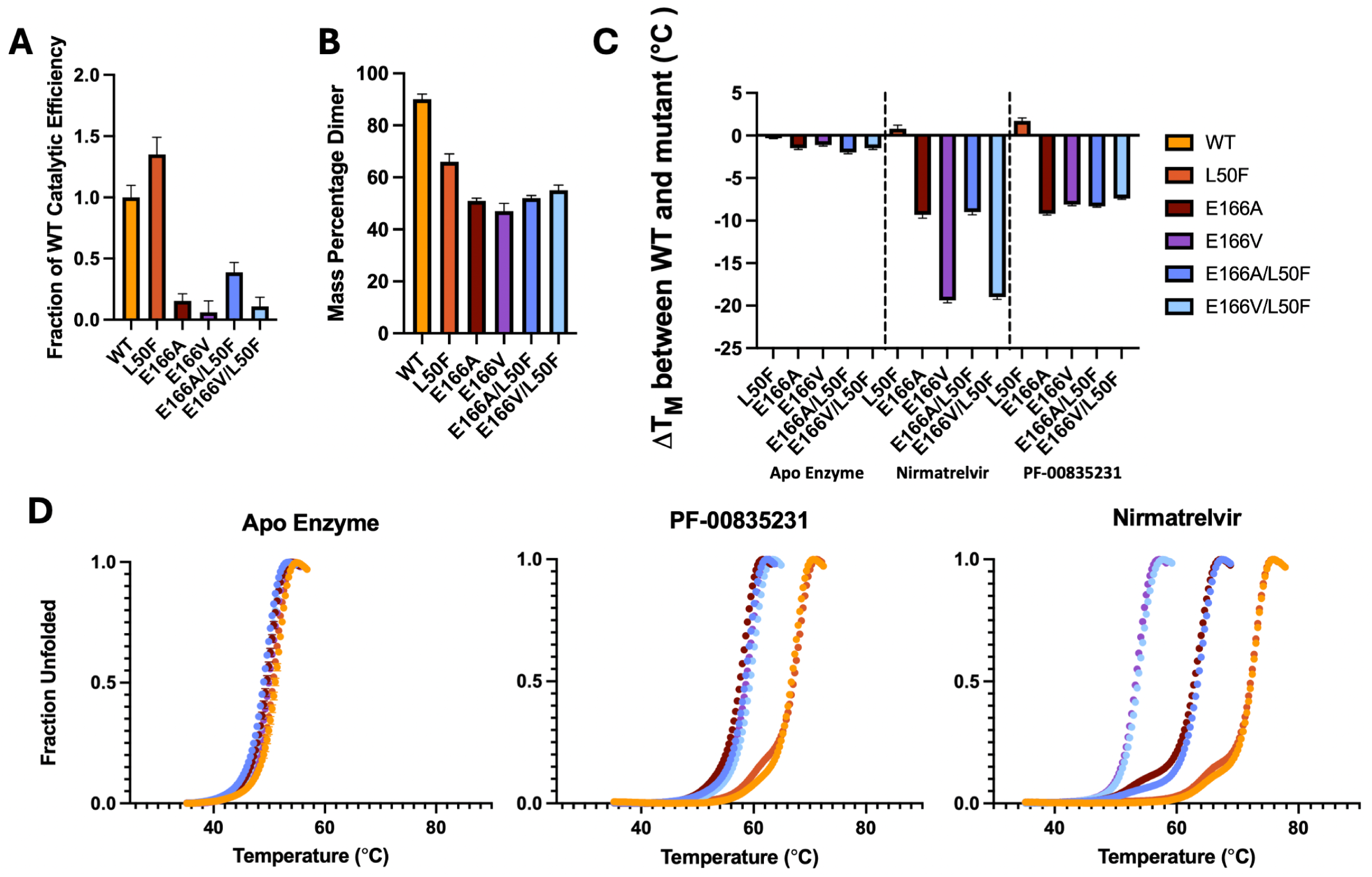
Enzymatic activity, dimerization, and stability of M^pro^ variants compared to WT enzyme. **A**) Fraction of catalytic efficiency (k_cat_/K_M_) relative to WT. **B**) The percentage of M^pro^ that is dimerized at a concentration of 5 μM detected by native mass-spectrometry **C**) Shift in the melting temperature (ΔT_m_) relative to WT enzyme in the apo and inhibitor-bound state for each variant. **D**) Melting curves for M^pro^ variants in the apo, PF-00835231 and nirmatrelvir bound state. The T_m_ was determined as the temperature where half of the enzyme was unfolded (see Methods).

**Table 2.** Enzyme kinetic parameters for E166 variants and WT M^pro^. Relative catalytic efficiency (k_cat_/K_M_) was calculated relative to WT.

| M<sup>pro</sup> Variant | $K_M$ (μM) | $V_{max}$ (μM/s) | $k_{cat}$ (1/s) | $k_{cat} / K_M$<br>1/(μMs) | Relative<br>$k_{cat} / K_M$ |
| --- | --- | --- | --- | --- | --- |
| WT | 23 ± 2 | 0.90 ± 0.03 | 12.0 ± 0.4 | 0.52 ± 0.05 | 100% |
| L50F | 20 ± 3 | 1.40 ± 0.06 | 13.8 ± 0.6 | 0.69 ± 0.10 | 133% |
| E166A | 47 ± 3 | 0.75 ± 0.02 | 3.75 ± 0.08 | 0.080 ± 0.005 | 15% |
| E166V | 42 ± 4 | 0.65 ± 0.02 | 1.30 ± 0.04 | 0.031 ± 0.003 | 6% |
| E166A/L50F | 37 ± 3 | 1.83 ± 0.06 | 7.33 ± 0.22 | 0.20 ± 0.02 | 38% |
| E166V/L50F | 46 ± 3 | 1.05 ± 0.03 | 2.63 ± 0.07 | 0.057 ± 0.005 | 11% |

### E166 variants have defective dimerization but WT-like thermostability

We utilized native mass spectrometry (nMS) to determine the fraction of dimeric versus monomeric M^pro^ in the absence of ligand. Measurements of dimerization were performed at 5 μM protein in ammonium acetate buffer (see Methods for details). WT M^pro^ at 5 μM was 90% dimerized (**Figure 2B**). Monomeric M^pro^ is known to be inactive^18^, and the monomer does not bind substrate with high affinity^19^. We have estimated the K_d_ for dimerization of WT M^pro^ to be 0.35 μM, which agrees with previous estimates from SAXS^18^, using a dilution nMS approach (**Figure S1**).

Dimerization in the absence of any ligand was quite reduced (47–66%; P<0.0001, n=3) in all E166 variants (**Figure 2B**). The L50F single-mutant variant also dimerized more poorly than WT enzyme (66–89%; P<0.0001, n=3). The double- and single-mutant E166 variants displayed similarly reduced dimerization relative to WT with no notable significant difference among them (P>0.01).

To assess the thermostability of the M^pro^ variants, we performed differential scanning fluorimetry (DSF) in the presence and absence of inhibitors. In the absence of inhibitors, the thermostabilities of these variants were roughly the same as WT M^pro^ (melting temperature T_M_ within 2 °C; **Figure 2C,D**). This indicates that the E166 variants are stable despite defunct dimerization.

Next, we tested the thermostability in the presence of saturating concentration of inhibitors (see Methods). The enzyme variants were incubated with excess inhibitor to ensure complete binding and ligand-induced dimerization (Supplemental Figure S2). As expected, inhibitor binding substantially increased the T_m_ of WT enzyme by 15–20 °C for both nirmatrelvir and PF- 00835231, indicating stabilization of WT M^pro^ by the bound inhibitor. In the presence of inhibitors, all variants were more thermostable than their comparable apo enzyme by at least 8°C. However, the E166 variants displayed reduced stability compared to WT enzyme. In the presence of PF-00835231, T_m_ was lower (by more than 7 °C) for the variants than that for WT (**Figure 2C,E**; **Table S1**). Against PF-00835231, these E166 variants had similar, low fold-changes in potency (<10-fold; **Table 1**) and the enzyme is expected to be fully inhibitor bound under the conditions tested. Analogous with their similar levels of drug resistance against PF- 00835231, the E166 variants had similar thermostabilities (T_m_ within 3 °C to one another) when bound to saturating amounts of PF-00835231. Unlike the E166 variants, the single-mutant variant L50F was an exception with a WT-like thermostability when bound to PF-00835231.

We found that drug-resistant E166 variants had highly deficient thermostabilities compared to WT when treated with a saturating concentration of nirmatrelvir (**Figure 2C; Figure S2, Table S1**). Considering nirmatrelvir resistance, the T_m_ of the four variants inversely correlates with the level potency loss we measured. The E166A variant, which shows a smaller 30-fold loss of nirmatrelvir potency relative to WT, yields a smaller 9.2 °C decrease in T_m_ compared to WT. The E166V variant has the largest amount of nirmatrelvir resistance (∼2700-fold) and accordingly shows a staggering 19.3 °C decrease in T_m_ compared to WT (**Figure 2C,F**;). All nirmatrelvir-bound enzymes were more thermostable than their comparable apo enzymes. The most resistant E166V and E166V/L50F had a T_m_ increase of a modest 3-4°C when nirmatrelvir-bound. In contrast, the less resistant variants had at least 13°C increase in their T_m_ when complexed with nirmatrelvir.

Alanine variants E166A and E166A/L50F show lower levels of nirmatrelvir resistance and lower ΔT_m_ whereas highly resistant valine variants E166V and E166V/L50F show high levels of nirmatrelvir fold-change in potency and a larger ΔT_m_. With PF-00835231, each variant we tested had similar, low levels of fold-change in potency (<10-fold) and accordingly, each variant had similar degrees of thermostability loss. For nirmatrelvir and PF-00835231 binding, our results suggest that fold-change in potency and inhibitor-bound thermostabilities are inversely correlated.

### Crystal structures of E166 M^pro^ variants reveal alternative conformation of the N-terminus

We determined the cocrystal structures of SARS-CoV-2 M^pro^ E166A, E166V, and E166V/L50F variants in complex with PF-00835231 to sub-1.9 Å resolution (**Table S2**). We performed cocrystallization trials with both nirmatrelvir and PF-00835231 for each of these M^pro^ variants; however, PF-00835231 complexes formed better-diffracting crystals with each variant.

Nirmatrelvir cocrystallization trials with these variants resulted in small, non-diffracting crystals. The E166A/L50F variant also failed to produce diffraction quality co-crystals with PF-00835231. The E166 variants co-crystallized as homodimers with two monomers in the asymmetric unit, in the P2_1_ space group. R-work for our structures ranges from 15-17% and R-free values range from 20-22% (**Table S2**). Both active sites were inhibitor-bound in every structure and the inhibitor occupied subsites S4 to S1’, as seen in our previous co-crystal structure of WT M^pro^ in complex with PF-00835231 (PDB ID: 8DSU). In each of our determined structures, PF- 00835231 was bound in the expected conformation^2^ with strong electron density observed for the covalent bond between Cys145 and the carbonyl of the hydroxymethylketone warhead of the inhibitor. In all the active sites, the carbinol of the hemithioketal is in a typical position oriented between catalytic residues His41 and Cys145.

SARS-CoV-2 M^pro^ substrate sequences universally feature a conserved glutamine residue at the P1 position. In WT M^pro^ complexed with either PF-00835231 or nirmatrelvir, which share identical P1 moieties, the inhibitor’s γ-lactam ring acts as a glutamine surrogate, mimicking substrate interactions with the protease. The inhibitor γ-lactam’s polar atoms hydrogen bond to Phe140 and His163 like in substrate binding^20^. In the active site of WT M^pro^, the S1 glutamine binding pocket is completed by the N-terminus connection with the loop spanning residues 160-170. In complex with PF-00835231 or nirmatrelvir, Glu166 makes three key hydrogen bonds to form the S1 subsite: between Glu166:OE2 and Ser1:OG; between Glu166:OE1 and Ser1:N; and between Glu166:OE1 and the nitrogen of the γ-lactam ring. However, in each of the inhibitor-bound Glu166 variant structures presented here, all three of these hydrogen bond interactions that define the S1 pocket are absent (**Figure 3**). These hydrogen bonding alterations, caused by mutations at residue 166, validate the critical role of E166 in completing and stabilizing the M^pro^ active site.

**Figure 3.**
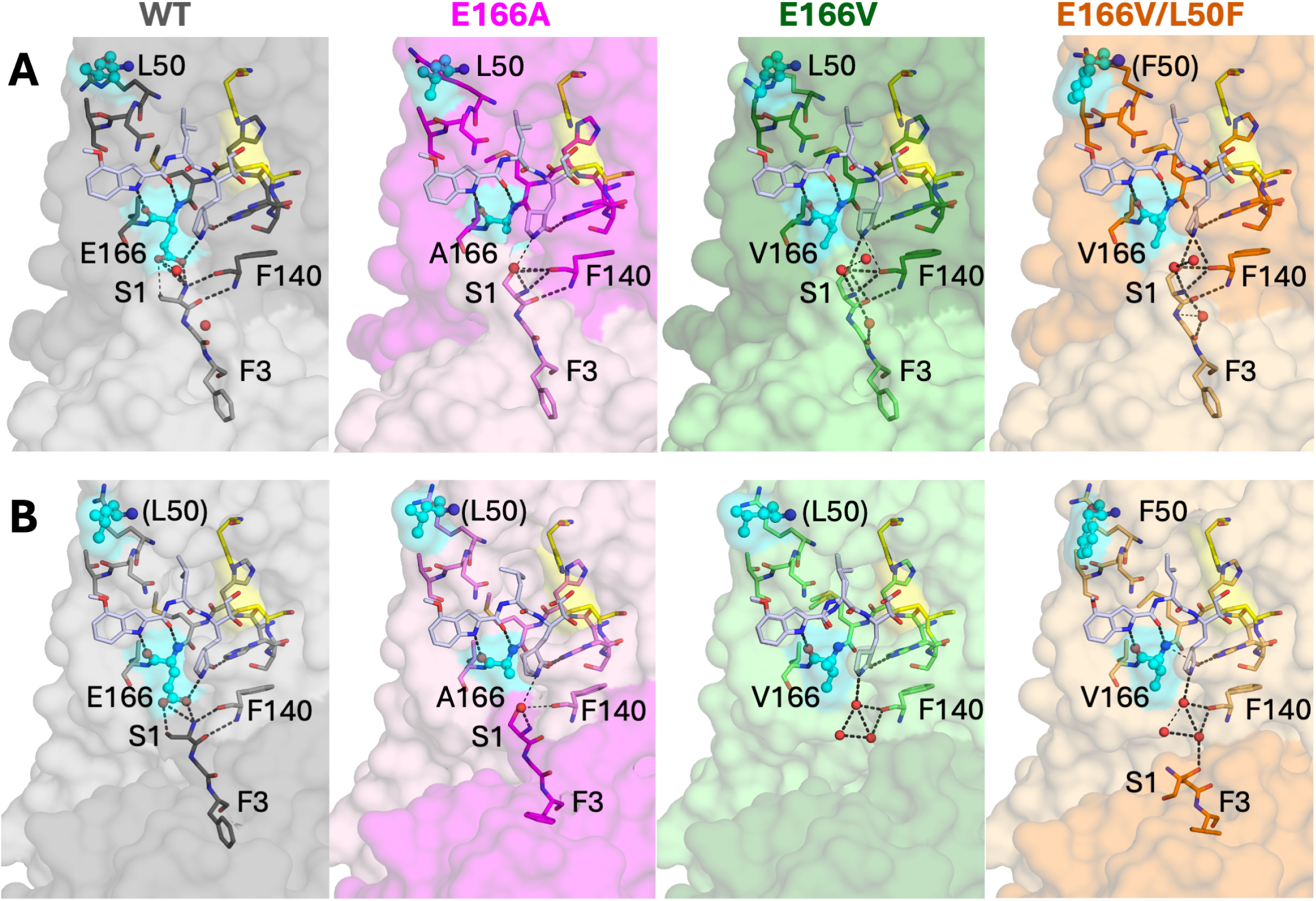
The active sites of the two protomers (**A** and **B**) in the cocrystal structures of M^pro^ variants with the inhibitor PF-00835231. The inhibitor is shown as blue-white sticks in each panel. The sites of mutations, residues 166 and 50, are highlighted as spheres and colored cyan. Monomers of each respective chain are distinguishable by color shade. Water molecules are shown as red spheres, and the hydrogen bonds are indicated by dashed lines.

In the WT PF-00835231-bound structure (PDB ID: 8DSU), the N-terminus is canonically positioned with Ser1 situated to form the S1 subsite. In contrast, these E166 M^pro^ variants show an alternative conformation of Ser1. Compared to the analogous PF-00835231-bound WT structure, the N-terminus of the Glu166 variants leans toward the inhibitor γ-lactam ring to fill the space normally occupied by Glu166 side-chain atoms (**Figure 3A**). Both E166A and E166V variant structures demonstrate this alternative N-terminus conformation. While the single-mutant variant L50F structure (PDB ID: 8E5C) retains WT-like positioning of the N-terminus, the double-mutant variant, E166V/L50F, displays the same alternative conformation of its N-terminus as the other two single-mutant variant E166 enzymes. The alternative conformation of the N-terminus, which has been discovered previously^13^, allows Ser1 to partially recapitulate the S1 site and complete the active site without the interactions from Glu166 side-chain atoms.

Our structures of E166A, E166V, and E166V/L50F M^pro^ variants demonstrate two N-terminus hydrogen bonds absent in the WT complex: between Ser1:OG hydrogen and Phe140:O and between Ser1:OG hydrogen and the nitrogen atom of the γ-lactam ring (**Figure 3A**). These hydrogen bonds are neither WT-like nor substrate mimicking. Despite the alternative N-terminus conformation, residues Gly2 and Phe3 retain WT-like positioning, and F140 maintains its WT-like conformation, forming direct hydrogen bonds to atoms of the γ-lactam ring.

In one monomer, the N-terminus is fully ordered in our electron density maps (**Figure 3A**), while in the other monomer, the N-terminal residues are either disordered, unresolved, or adopt an alternative conformation (**Figure 3B**). The differences highlight how E166 mutations might be disrupting the local structure of the active site. In the E166A variant structure, both active sites showed a resolved N-terminus in the same non-canonical position. The less ordered electron density in one monomer causes the appearance of an incomplete S1 pocket for E166V, with residues Ser1 and Gly2 unresolved (**Figure 3B**). The N-terminus of E166V/L50F was well-resolved in an atypical position, folded away from the active site (**Figure 3B**). One other crystal structure of E166V M^pro^ bound to ensitrelvir demonstrated a similar conformation wherein the N-terminus was folded far away from the S1 site at only one monomer of the dimer^13^.

## Discussion

In this study, we aimed to investigate how mutations at the E166 residue in SARS-CoV-2 M^pro^ impact drug resistance and enzymatic function, focusing on E166A and E166V variants with or without the addition of L50F, a compensatory mutation. These variants demonstrated reduced potencies of antiviral drug nirmatrelvir and the analog PF-00835231 at the cost of significantly compromised catalytic efficiency. Resistance against nirmatrelvir was modest to high, but resistance to PF-00835231 was relatively limited. The distal L50F mutation exhibited a compensatory effect, partially restoring the loss in enzymatic activity for drug resistant M^pro^ variants. Our results highlight the critical role of E166 in stabilizing inhibitor interactions and fully forming the active site, supported by apo versus inhibitor-bound thermostability and dimerization measurements. Our cocrystal structures of E166A, E166V, and E166V/L50F M^pro^ variants in complex with PF-00835231 revealed that interactions between the S1 pocket and the E166 side chain are essential for both active site completion and dimerization.

We have shown that the addition of L50F to drug resistant SARS-CoV-2 M^pro^ E166 variants compensates for defective enzymatic activities. The compensatory mutation L50F has been discovered *in vitro*^10,11,16^ and by viral passaging^3–5^; this mutation on its own enhances enzymatic activity relative to WT M^pro^ resulting in hyperactivity^16^. Our results agree with this effect of L50F, as we found that this mutation partially restores the catalytic efficiency in drug resistant, activity-compromised variants E166A and E166V. This compensation is a crucial mechanism in enabling drug resistance to emerge, since drug resistant enzymes often show compromised activity that critically diminishes viral fitness. Furthermore, the L50 position of M^pro^ has been subject to numerous single amino acid substitutions and/or deletions in its evolutionary history across divergent genera of coronaviruses (**Figure S3**). This variability highlights the adaptability of coronavirus M^pro^ suggesting that the residue substitutions at this position may serve as adaptive pathways for these viruses to navigate antiviral selective pressures. Understanding the role of L50F in rescuing or enhancing enzymatic activity may be critical for developing strategies to combat emerging drug resistance in coronaviruses.

Drug resistance mutations at E166 in M^pro^ significantly reduce affinity for nirmatrelvir but preserve substrate cleavage. For example, the E166V mutation reduced nirmatrelvir potency by about 2700-fold but the catalytic efficiency was decreased by only 2-fold. This discrepancy highlights a crucial aspect of E166 drug resistance: the mutations specifically disrupt interactions critical for the inhibitor without severely impairing substrate binding or processing. Previously, our comparison of the substrate envelope with the crystal structures of WT M^pro^ in complex with PF-00835231 and nirmatrelvir indicated that residue E166 is among those vulnerable to drug resistance mutations^20^. Both inhibitors protrude from the substrate envelope at the P3 and P4 sites and interact with E166. These contacts made outside of the substrate envelope render E166 a key potential site for resistance against both inhibitors, but the relative flexibility of PF-00835231 might allow better accommodation of E166 mutations. The mutability of E166 under selective pressure emphasizes the need for therapeutic strategies that avoid such vulnerability and address the adaptive elasticity of viral proteases.

We hypothesize that structural waters may serve a key function in the more ordered and well-resolved active sites of drug resistant M^pro^ E166 variants. Certain structural waters differ between WT and drug-resistant M^pro^ variants. In the E166V and E166V/L50F variant structures, water molecules are positioned to substitute for the OE1 atom of Glu166, facilitating interactions with Phe140 and the inhibitor. A water molecule also recreates the hydrogen bond between Ser1 and Phe140, stabilizing Phe140’s position, which is critical for binding substrate and inhibitors. These conserved structural waters suggest that the S1 site can be transiently completed by solvent molecules in the absence of side chain polar atoms from Glu166, allowing E166 variants to transiently bind and cleave substrates while being deficient in stable inhibitor binding. For nirmatrelvir binding, the transient substitution of E166 atoms with water molecules may contribute to the variants’ increased drug resistance and our inability to cocrystallize them with established nirmatrelvir cocrystallization conditions. We suppose based on our cocrystal structures that the interactions lost from the missing E166 atoms can be somewhat rescued by structural waters, which may be essential for E166A and E166V variants to complete a functional M^pro^ active site, to bind ligands, or to stably dimerize.

SARS-CoV-2 remains a healthcare concern with the potential emergence of drug resistance in M^pro^, and E166 is a key site of mutation under the selection pressure of nirmatrelvir. Drug resistance mutations at E166, such as E166A and E166V, significantly reduce inhibitor potency while preserving substrate binding, making this conserved residue a focal point for resistance studies^4,6,7,10,11^. However, these variants have severely reduced catalytic efficiency, which would impact viral fitness in agreement with their absence from clinical samples. Despite its conservation across all coronavirus clades (**Figure S3**), mutational scanning shows that Glu at the 166 position is not necessarily essential^8^, highlighting its mutability and susceptibility to resistance mutations. Zoonotic transmission events documented in recent history highlight that coronaviruses such as SARS-CoV-2 and MERS-CoV can be transmitted from animals to humans with significant health consequences, and as viral spillovers continue, future coronavirus outbreaks are likely. Given E166’s shared vulnerability across coronaviral strains, resistant variants observed in SARS-CoV-2, including E166A and E166V, could similarly arise in newly emergent coronaviruses—particularly since nirmatrelvir would likely be repurposed as a first-line antiviral option.

Compensatory mutations, such as L50F, further complicate the resistance landscape by restoring catalytic efficiency in E166 variants. The double-mutant variants E166A/L50F and E166V/L50F investigated here revealed that compensatory mutations can mitigate the functional deficits caused by primary resistance mutations. Understanding viral adaptive mechanisms is critical for designing inhibitors that counteract both resistance and compensation, offering strategies to combat current and future coronavirus threats.

## Methods

### Protein purification

Wild-type and variant M^pro^ proteins were expressed with an N-terminal polyhistidine-SUMO tag and purified as described previously^9,16,20,21^. The point mutations were introduced into the pETite expression plasmid using site-directed mutagenesis.

### Crystallization

For cocrystallization, complexes with PF-00835231 were assembled by incubating 6 mg/mL of each M^pro^ with a 10-fold molar excess of inhibitor for 1 h at room temperature. The solutions were spun down at 10,000xg for 10 minutes to remove any insoluble compound or protein aggregates. Protein crystals were obtained with 10–20% (*w*/*v*) PEG 3350, 0.2 M NaCl, and 0.1 M Bis-Tris Methane pH 5.5 by hanging drop vapor diffusion at room-temperature in pre-greased VDX trays (Hampton Research, Aliso Viejo, CA, USA). Varying the protein-to-mother liquor ratios (1 µL:2 µL, 2 µL:2 µL, 3 µL:2 µL) helped obtain large, diffraction-quality crystals. To limit vibrational stress, crystallization trays were placed on foam padding. Cocrystals of M^pro^ variants with PF-00835231 appeared overnight and grew fully within 2 weeks.

### Data collection and structure determination

Crystals were sent for data collection at the Brookhaven National Laboratory NSLS-II Beamline 17-ID-2 (FMX). X-ray diffraction data were collected at 100 K. Cocrystals were soaked in cryogenic solutions made by supplementing the exact precipitant solutions with 25% glycerol, then looped and frozen in liquid nitrogen. At NSLS-II, the collected diffraction intensities were automatically indexed, integrated, and scaled using XDS^22^. Prior to molecular replacement, the model was modified by removing all water, buffer, and cryogenic molecules as well as the small molecule inhibitor in the active site. All structures were determined using molecular replacement with PHASER^23^. The reference model used was PDB ID: 8DSU. To minimize reference model bias, 5% of the data was reserved to calculate R ^24^. Before fitting inhibitor atoms into the electron density, inhibitor geometry was optimized in Gaussview 6 using Gaussian 16 with the basis set: DFT B3LYP 6-311++G (d,p). Model building and refinement were performed using Coot^25^ and Phenix^26^. X-ray data collection parameters and refinement statistics are presented in Supplemental Material, Table S2.

### Structural analysis

Each of the cocrystal structures contained an M^pro^ dimer in the asymmetric unit. All structural figures were generated and analyses performed in PyMOL by Schrödinger, LLC^27^. Key residues which are stubbed in the cocrystal structures were added to the structural figures; the side- chains of key stubbed residues are shown, but in these cases the residue label is placed within parentheses, e.g. (F50). Hydrogen bonds were determined using the show_contacts PyMOL plugin with default parameters where the bond angle is between 63 and 180 degrees and the distance less than 4.0 Å for any and 3.6 Å for an ideal hydrogen bond between the proton and heavy atom.

### Differential scanning fluorimetry (DSF)

Thermal shift assays were performed in final conditions of 2 µM M^pro^, 50 mM Tris pH 8.0, 300 mM NaCl, 2% DMSO, with and without each inhibitor, and with 5x Sypro orange dye (Invitrogen 5000x Sypro). Initial experiments with WT M^pro^ demonstrated that 2 µM of M^pro^ produces optimal signal:noise ratio under our experimental conditions. M^pro^ variants were mixed with inhibitor and incubated at room temperature for an hour. Inhibitor concentration was determined to ensure complete saturation of all protease variants according to the potency against the most resistant M^pro^ variant; for example, E166V requires 400 µM of nirmatrelvir to reach 0% activity, so 400 µM nirmatrelvir was used for all M^pro^ variants in the assay. To achieve apo protease melting curves, DMSO was added to 2% in wells with no inhibitor. After 1 hour incubation, 5000x Sypro orange dye was diluted to a final concentration of 5x on the plate, and was added only immediately before the assay began. Thermal denaturation spectra were run in a Thermo Scientific AB-0700 96-well OCR plate using a Bio-Rad CFX Real Time PCR thermocycler. Temperatures during the melt ranged from 25 – 95 °C, with intervals of 0.3 °C every 12 seconds. The HEX channel was utilized to measure relative fluorescence units (RFU) for each well at each temperature.

Data were normalized to values between 0 and 1 using the normalization equation: F_norm_ = F - F_min_ / (F_max_ – F_min_). Values were then truncated after three data points past the maximum value of 1. A Boltzmann sigmoidal model was fit to the normalized data to calculate the melting temperature (*T_m_*). The equation Δ*T*_m_ = *T*_m (inhibitor)_ − *T*_m (DMSO)_ was used to calculate the thermal shift (Δ*T*_m_). The *T_m_* values are presented as mean ± standard error (12 data points for each of the 3 replicates). The calculated *T_m_* values were confirmed also by plotting the first derivative (*d*RFU/*d*T) vs temperature to determine the temperature at which *d*RFU/*d*T is maximum — the *T_m_*. These values matched very closely (within ± 0.3 °C) with the *T_m_* from Boltzmann fits.

### Enzyme activity assays

To determine the enzyme kinetic parameters of M^pro^ variants, 75 to 500 nM enzyme was added to a series of 0–200 μM FRET substrate (Dabcyl-KTSAVLQSGFRKM-Glu(Edans) (GenScript)) in assay buffer (50 mM Tris pH 7.5, 50 mM NaCl, 1 mM ethylenediaminetetraacetic acid (EDTA), 1 mM dithiothreitol (DTT), and 4% dimethyl sulfoxide (DMSO)). The cleavage reaction was monitored using a PerkinElmer Envision plate reader at room-temperature (340 nm excitation and 492 nm emission). Three or more replicates were performed for each variant. The initial velocities (RFU/s) were plotted against substrate concentrations and fit using GraphPad Prism 10^28^ to the Michaelis–Menten equation. RFU was converted to concentration using a linear fit of the calibration curve obtained with different concentrations of substrate and the end point of cleavage product with WT enzyme, which gave a factor of 6000 RFU per μM under the assay conditions.

### Enzyme inhibition assays

Inhibition assays were performed in the same assay buffer (50 mM Tris pH 7.5, 50 mM NaCl, 1 mM ethylenediaminetetraacetic acid (EDTA), 1 mM dithiothreitol (DTT), but with only 1% DMSO. To determine the inhibition constant *K_i_*, enzyme was incubated at room temperature with increasing concentrations of nirmatrelvir or PF-00835231 for 1 hour in assay buffer. The enzymatic reaction was initiated with 40 μM protease FRET substrate and monitored using a PerkinElmer Envision plate reader. At least three replicates were performed for each inhibitor concentration. The initial velocity for each reaction was calculated by linear regression.

The *K_i_* was calculated by plotting the initial velocity (RFU/s) at each inhibitor concentration (µM) and then fit to the Morrison equation, using each enzyme’s respective K_M_, in GraphPad Prism 10^28^ software. Enzyme levels were adjusted according to the K_i_ estimates from initial experiments to maintain [E]/K_i_<100 to ensure confidence in K_i_ determinations. In cases of very potent inhibition where the enzyme concentration could not be decreased further due to loss of signal, [E]/100 is reported as the upper estimate for K_i_ as the assay detection limit.

### Native Mass Spectrometry

Each protein was purified in SEC buffer (25 mM HEPES pH 7.5, 150 mM NaCl, and 1 mM TCEP) and then diluted to 2-4 mg/mL in 50-100 µL in the same buffer. The protein was dialyzed for three hours into 200 mM ammonium acetate (pH 6.8) using a Slide-A-Lyzer MINI Dialysis Device (0.5 mL; MWCO 20 kDa; Thermo Fisher Scientific), then dialyzed again overnight. After dialysis, the protein concentration was re-determined based on absorbance at 280 nm. Each protein was diluted into ammonium acetate to 5 µM. Native mass-spectrometry analysis was performed on a QE-UHMR mass spectrometer (Advion Interchim Scientific) with parameters as follows: Positive mode; spray voltage: 1.5 kV; resolution: 12,500; mass range (*m*/z): 2000-8000. The mass spectra were analyzed using PMI-Byos Intact software (Protein Metrics Inc.). Each protein was analyzed in triplicate.

## Data availability

The data that support this study are available from the corresponding authors upon reasonable request. The crystal structures determined in the current study are available in the Protein Data Bank (https://www.rcsb.org) with accession codes 9EL4, 9ELV, and 9MEI.

## References

(1) Boras, B.; Jones, R. M.; Anson, B. J.; Arenson, D.; Aschenbrenner, L.; Bakowski, M. A.;Beutler, N.; Binder, J.; Chen, E.; Eng, H.; Hammond, H.; Hammond, J.; Haupt, R. E.; Hoffman, R.; Kadar, E. P.; Kania, R.; Kimoto, E.; Kirkpatrick, M. G.; Lanyon, L.; Lendy, E. K.; Lillis, J. R.; Logue, J.; Luthra, S. A.; Ma, C.; Mason, S. W.; McGrath, M. E.; Noell, S.;Obach, R. S.; O’ Brien, M. N.; O’Connor, R.; Ogilvie, K.; Owen, D.; Pettersson, M.; Reese, M. R.; Rogers, T. F.; Rosales, R.; Rossulek, M. I.; Sathish, J. G.; Shirai, N.; Steppan, C.; Ticehurst, M.; Updyke, L. W.; Weston, S.; Zhu, Y.; White, K. M.; García-Sastre, A.; Wang, J.; Chatterjee, A. K.; Mesecar, A. D.; Frieman, M. B.; Anderson, A. S.; Allerton, C. Preclinical Characterization of an Intravenous Coronavirus 3CL Protease Inhibitor for the Potential Treatment of COVID19. Nat Commun 2021, 12 (1), 6055. 10.1038/s41467-021-26239-2.

(2) Hoffman, R. L.; Kania, R. S.; Brothers, M. A.; Davies, J. F.; Ferre, R. A.; Gajiwala, K. S.; He, M.; Hogan, R. J.; Kozminski, K.; Li, L. Y.; Lockner, J. W.; Lou, J.; Marra, M. T.; Mitchell, L. J.; Murray, B. W.; Nieman, J. A.; Noell, S.; Planken, S. P.; Rowe, T.; Ryan, K.; Smith, G. J.; Solowiej, J. E.; Steppan, C. M.; Taggart, B. Discovery of Ketone-Based Covalent Inhibitors of Coronavirus 3CL Proteases for the Potential Therapeutic Treatment of COVID-19. J Med Chem 2020, 63 (21), 12725–12747. 10.1021/acs.jmedchem.0c01063.

(3) Zhou, Y.; Gammeltoft, K. A.; Ryberg, L. A.; Pham, L. V.; Tjørnelund, H. D.; Binderup, A.; Duarte Hernandez, C. R.; Fernandez-Antunez, C.; Offersgaard, A.; Fahnøe, U.; Peters, G. H. J.; Ramirez, S.; Bukh, J.; Gottwein, J. M. Nirmatrelvir-Resistant SARS-CoV-2 Variants with High Fitness in an Infectious Cell Culture System. Sci Adv 2022, 8 (51), eadd7197. 10.1126/sciadv.add7197.

(4) Iketani, S.; Mohri, H.; Culbertson, B.; Hong, S. J.; Duan, Y.; Luck, M. I.; Annavajhala, M. K.; Guo, Y.; Sheng, Z.; Uhlemann, A.-C.; Goff, S. P.; Sabo, Y.; Yang, H.; Chavez, A.; Ho, D. D. Multiple Pathways for SARS-CoV-2 Resistance to Nirmatrelvir. Nature 2023, 613 (7944), 558–564. 10.1038/s41586-022-05514-2.

(5) Zhu, Y.; Yurgelonis, I.; Noell, S.; Yang, Q.; Guan, S.; Li, Z.; Hao, L.; Rothan, H.; Rai, D. K.; McMonagle, P.; Baniecki, M. L.; Greasley, S. E.; Plotnikova, O.; Lee, J.; Nicki, J. A.; Ferre, R.; Byrnes, L. J.; Liu, W.; Craig, T. K.; Steppan, C. M.; Liberator, P.; Soares, H. D.; Allerton, C. M. N.; Anderson, A. S.; Cardin, R. D. In Vitro Selection and Analysis of SARS-CoV-2 Nirmatrelvir Resistance Mutations Contributing to Clinical Virus Resistance Surveillance. Sci. Adv. 2024, 10 (30), eadl4013. 10.1126/sciadv.adl4013.

(6) Hu, Y.; Lewandowski, E. M.; Tan, H.; Zhang, X.; Morgan, R. T.; Zhang, X.; Jacobs, L. M. C.; Butler, S. G.; Gongora, M. V.; Choy, J.; Deng, X.; Chen, Y.; Wang, J. Naturally Occurring Mutations of SARS-CoV-2 Main Protease Confer Drug Resistance to Nirmatrelvir. ACS Cent. Sci. 2023, 9 (8), 1658–1669. 10.1021/acscentsci.3c00538.

(7) Padhi, A. K.; Tripathi, T. Hotspot Residues and Resistance Mutations in the Nirmatrelvir-Binding Site of SARS-CoV-2 Main Protease: Design, Identification, and Correlation with Globally Circulating Viral Genomes. Biochem Biophys Res Commun 2022, 629, 54–60. 10.1016/j.bbrc.2022.09.010.

(8) Flynn, J. M.; Samant, N.; Schneider-Nachum, G.; Barkan, D. T.; Yilmaz, N. K.; Schiffer, C. A.; Moquin, S. A.; Dovala, D.; Bolon, D. N. A. Comprehensive Fitness Landscape of SARS-CoV-2 Mpro Reveals Insights into Viral Resistance Mechanisms. Elife 2022, 11, e77433. 10.7554/eLife.77433.

(9) Flynn, J. M.; Huang, Q. Y. J.; Zvornicanin, S. N.; Schneider-Nachum, G.; Shaqra, A. M.; Yilmaz, N. K.; Moquin, S. A.; Dovala, D.; Schiffer, C. A.; Bolon, D. N. A. Systematic Analyses of the Resistance Potential of Drugs Targeting SARS-CoV-2 Main Protease. ACS Infect Dis 2023, 9 (7), 1372–1386. 10.1021/acsinfecdis.3c00125.

(10) Jochmans, D.; Liu, C.; Donckers, K.; Stoycheva, A.; Boland, S.; Stevens, S. K.; De Vita, C.; Vanmechelen, B.; Maes, P.; Trüeb, B.; Ebert, N.; Thiel, V.; De Jonghe, S.; Vangeel, L.; Bardiot, D.; Jekle, A.; Blatt, L. M.; Beigelman, L.; Symons, J. A.; Raboisson, P.; Chaltin, P.; Marchand, A.; Neyts, J.; Deval, J.; Vandyck, K. The Substitutions L50F, E166A, and L167F in SARS-CoV-2 3CLpro Are Selected by a Protease Inhibitor In Vitro and Confer Resistance To Nirmatrelvir. mBio 2023, 14 (1), e0281522. 10.1128/mbio.02815-22.

(11) Gammeltoft, K. A.; Zhou, Y.; Ryberg, L. A.; Pham, L. V.; Binderup, A.; Hernandez, C. R. D.; Offersgaard, A.; Fahnøe, U.; Peters, G. H. J.; Ramirez, S.; Bukh, J.; Gottwein, J. M. Substitutions in SARS-CoV-2 Mpro Selected by Protease Inhibitor Boceprevir Confer Resistance to Nirmatrelvir. Viruses 2023, 15 (9), 1970. 10.3390/v15091970.

(12) Noske, G. D.; de Souza Silva, E.; de Godoy, M. O.; Dolci, I.; Fernandes, R. S.; Guido, R. V. C.; Sjö, P.; Oliva, G.; Godoy, A. S. Structural Basis of Nirmatrelvir and Ensitrelvir Activity against Naturally Occurring Polymorphisms of the SARS-CoV-2 Main Protease. J Biol Chem 2023, 299 (3), 103004. 10.1016/j.jbc.2023.103004.

(13) Duan, Y.; Zhou, H.; Liu, X.; Iketani, S.; Lin, M.; Zhang, X.; Bian, Q.; Wang, H.; Sun, H.; Hong, S. J.; Culbertson, B.; Mohri, H.; Luck, M. I.; Zhu, Y.; Liu, X.; Lu, Y.; Yang, X.; Yang, K.; Sabo, Y.; Chavez, A.; Goff, S. P.; Rao, Z.; Ho, D. D.; Yang, H. Molecular Mechanisms of SARS-CoV-2 Resistance to Nirmatrelvir. Nature 2023, 622 (7982), 376–382. 10.1038/s41586-023-06609-0.

(14) Havranek, B.; Demissie, R.; Lee, H.; Lan, S.; Zhang, H.; Sarafianos, S.; Ayitou, A. J.-L.; Islam, S. M. Discovery of Nirmatrelvir Resistance Mutations in SARS-CoV-2 3CLpro: A Computational-Experimental Approach. J Chem Inf Model 2023, 63 (22), 7180–7188. 10.1021/acs.jcim.3c01269.

(15) Henes, M.; Lockbaum, G. J.; Kosovrasti, K.; Leidner, F.; Nachum, G. S.; Nalivaika, E. A.; Lee, S.-K.; Spielvogel, E.; Zhou, S.; Swanstrom, R.; Bolon, D. N. A.; Kurt Yilmaz, N.; Schiffer, C. A. Picomolar to Micromolar: Elucidating the Role of Distal Mutations in HIV-1 Protease in Conferring Drug Resistance. ACS Chem Biol 2019, 14 (11), 2441–2452. 10.1021/acschembio.9b00370.

(16) Flynn, J. M.; Zvornicanin, S. N.; Tsepal, T.; Shaqra, A. M.; Kurt Yilmaz, N.; Jia, W.; Moquin, S.; Dovala, D.; Schiffer, C. A.; Bolon, D. N. A. Contributions of Hyperactive Mutations in Mpro from SARS-CoV-2 to Drug Resistance. ACS Infect Dis 2024, 10 (4), 1174–1184. 10.1021/acsinfecdis.3c00560.

(17) Zhang, L.; Xie, X.; Luo, H.; Qian, R.; Yang, Y.; Yu, H.; Huang, J.; Shi, P.-Y.; Hu, Q. Resistance Mechanisms of SARS-CoV-2 3CLpro to the Non-Covalent Inhibitor WU-04. Cell Discov 2024, 10 (1), 40. 10.1038/s41421-024-00673-0.

(18) Silvestrini, L.; Belhaj, N.; Comez, L.; Gerelli, Y.; Lauria, A.; Libera, V.; Mariani, P.; Marzullo, P.; Ortore, M. G.; Palumbo Piccionello, A.; Petrillo, C.; Savini, L.; Paciaroni, A.; Spinozzi, F. The Dimer-Monomer Equilibrium of SARS-CoV-2 Main Protease Is Affected by Small Molecule Inhibitors. Sci Rep 2021, 11 (1), 9283. 10.1038/s41598-021-88630-9.

(19) El-Baba, T. J.; Lutomski, C. A.; Kantsadi, A. L.; Malla, T. R.; John, T.; Mikhailov, V.; Bolla, J. R.; Schofield, C. J.; Zitzmann, N.; Vakonakis, I.; Robinson, C. V. Allosteric Inhibition of the SARS-CoV-2 Main Protease: Insights from Mass Spectrometry Based Assays*. Angew Chem Int Ed Engl 2020, 59 (52), 23544–23548. 10.1002/anie.202010316.

(20) Shaqra, A. M.; Zvornicanin, S. N.; Huang, Q. Y. J.; Lockbaum, G. J.; Knapp, M.; Tandeske, L.; Bakan, D. T.; Flynn, J.; Bolon, D. N. A.; Moquin, S.; Dovala, D.; Kurt Yilmaz, N.; Schiffer, C. A. Defining the Substrate Envelope of SARS-CoV-2 Main Protease to Predict and Avoid Drug Resistance. Nat Commun 2022, 13 (1), 3556. 10.1038/s41467-022-31210-w.

(21) Zvornicanin, S. N.; Shaqra, A. M.; Huang, Q. J.; Ornelas, E.; Moghe, M.; Knapp, M.; Moquin, S.; Dovala, D.; Schiffer, C. A.; Kurt Yilmaz, N. Crystal Structures of Inhibitor-Bound Main Protease from Delta- and Gamma-Coronaviruses. Viruses 2023, 15 (3), 781. 10.3390/v15030781.

(22) Kabsch, W. XDS. Acta Crystallogr D Biol Crystallogr 2010, 66 (Pt 2), 125–132. 10.1107/S0907444909047337.

(23) McCoy, A. J.; Grosse-Kunstleve, R. W.; Adams, P. D.; Winn, M. D.; Storoni, L. C.; Read, R. J. Phaser Crystallographic Software. J Appl Crystallogr 2007, 40 (Pt 4), 658–674. 10.1107/S0021889807021206.

(24) Brünger, A. T. Free R Value: A Novel Statistical Quantity for Assessing the Accuracy of Crystal Structures. Nature 1992, 355 (6359), 472–475. 10.1038/355472a0.

(25) Emsley, P.; Cowtan, K. Coot: Model-Building Tools for Molecular Graphics. Acta Crystallogr D Biol Crystallogr 2004, 60 (Pt 12 Pt 1), 2126–2132. 10.1107/S0907444904019158.

(26) Adams, P. D.; Afonine, P. V.; Bunkóczi, G.; Chen, V. B.; Davis, I. W.; Echols, N.; Headd, J. J.; Hung, L.-W.; Kapral, G. J.; Grosse-Kunstleve, R. W.; McCoy, A. J.; Moriarty, N. W.; Oeffner, R.; Read, R. J.; Richardson, D. C.; Richardson, J. S.; Terwilliger, T. C.; Zwart, P. H. PHENIX: A Comprehensive Python-Based System for Macromolecular Structure Solution. Acta Crystallogr D Biol Crystallogr 2010, 66 (Pt 2), 213–221. 10.1107/S0907444909052925.

(27) The PyMOL Molecular Graphics System.

